# Context and Domain Matter: The Error-Related Negativity in Peer Presence Predicts Fear of Negative Evaluation, not Global Social Anxiety, in Adolescents

**DOI:** 10.1101/2022.07.01.498524

**Authors:** Yanbin Niu, Zixuan Li, Jeremy W. Pettit, George A. Buzzell, Jingjing Zhao

**Author notes:** **Co-corresponding/Senior Authors:** Jingjing Zhao, Ph.D., School of Psychology, Shaanxi Normal University, 199 South Chang’an Road, Xian, Shaanxi, China 710062, George A. Buzzell, Ph.D., Florida International University and the Center for Children and Families, 11200 SW 8^th^ Street, DM 256, Miami, FL, USA 33199. These authors contributed equally to this manuscript (co-corresponding/senior authors) (George A. Buzzell, Jingjing Zhao).

## Abstract

**Background:** Social anxiety symptoms are most likely to emerge during adolescence, a developmental window marked by heightened concern over peer evaluation. However, the neurocognitive mechanism(s) underlying adolescent social anxiety remain unclear. Emerging work points to the error-related negativity (ERN) as a potential neural marker of exaggerated self/error-monitoring in social anxiety, particularly for errors committed in front of peers.However, social anxiety symptoms are marked by heterogeneity and it remains unclear exactly what domain(s) of social anxiety symptoms are associated with ERN variation in peer presence, particularly within the adolescent period.

**Methods:** To advance and deepen the mechanistic understanding of the ERN’s putative role as a neural marker for social anxiety in adolescence, we leveraged a social manipulation procedure and assessed a developmentally-salient domain of social anxiety during adolescence—Fear of Negative Evaluation (FNE). Adolescents residing in Hanzhong, a small city in the southwestern region of mainland China, had EEG recorded while performing a flanker task, twice (peer presence/absence); FNE, as well as global social anxiety symptoms were assessed.

**Results:** Overall ERN increases in peer presence. FNE specifically, but not global levels of social anxiety symptoms, predicted ERN in peer presence.

**Conclusions:** These data are the first demonstration that the ERN relates to a specific domain of social anxiety in adolescents, as well as the first evidence of such relations within a non-WEIRD (Western, Educated, Industrialized, Rich and Democratic) sample. Results have important implications for theory and research into adolescent social anxiety.

---

Social anxiety refers to a cluster of symptoms, including an intense fear of social situations or social scrutiny (Morrison & Heimberg, 2013; Rapee & Heimberg, 1997). These symptoms are most likely to emerge during adolescence (Kessler et al., 2005), a developmental window marked by increased social motivation and heightened concern over peer evaluation (Crone & Dahl, 2012; Dahl, 2004; Parker, Rubin, Erath, Wojslawowicz, & Buskirk, 2006; Steinberg & Morris, 2001). Cognitive models suggest that social anxiety involves negative biases in cognitive processing, including biases toward internal sources of social threat (committing errors/mistakes, autonomic responses), and greater self-monitoring within social settings, among other cognitive processes (Clark & Wells, 1995; Pineles & Mineka, 2005; Rapee & Heimberg, 1997). To extend cognitive models and advance a mechanistic understanding of social anxiety, it is important to identify neurocognitive processes linked to social anxiety symptoms in adolescence. In the current study, we focus on the error-related negativity (ERN) as a theoretically-relevant neurocognitive process.

Prior work demonstrates that the ERN, an event-related potential (ERP) elicited by errors of commission (Falkenstein, Hohnsbein, Hoormann, & Blanke, 1991; Gehring, Goss, Coles, Meyer, & Donchin, 1993), can serve as a neural marker of exaggerated error monitoring (self-monitoring) in individuals with general anxiety (Hajcak, Klawohn, & Meyer, 2019; Meyer, 2022; Moser, Moran, Schroder, Donnellan, & Yeung, 2013; Olvet & Hajcak, 2008) and social anxiety (Endrass, Riesel, Kathmann, & Buhlmann, 2014; Kujawa et al., 2016; Meyer, Carlton, Crisler, & Kallen, 2018). Moreover, the ERN predicts a pattern of error-related autonomic responses that are typically observed in response to threat (Hajcak et al., 2003; Hajcak & Foti, 2008). In addition to the ERN, the correct related negativity (CRN) is triggered by correct responses with a similar latency in the response-locked waveform as the ERN but with a smaller amplitude (Ford, 1999; Vidal et al., 2000, 2003). The CRN appears to have similar morphological and topographical properties to ERN, and is thought to reflect self-monitoring on correct trials (Allain et al., 2004; Suchan et al., 2007; Vidal et al., 2000).

Emerging work investigates associations between social anxiety and ERN specifically when errors are committed in social (vs. nonsocial) settings (Barker, Troller-Renfree, Pine, & Fox, 2015, 2015; Buzzell et al., 2017b; Voegler et al., 2018). Yet such work has yielded mixed results in terms of the link between global social anxiety symptoms and the ERN, and only one study has investigated these relations during adolescence (Buzzell, Troller-Renfree, et al., 2017). Given that social anxiety symptomology is marked by considerable heterogeneity (Hofmann, Heinrichs, & Moscovitch, 2004; Spokas & Cardaciotto, 2014; Yu, Zhou, Wang, & Tang, 2020), inconsistent associations between the ERN and social anxiety could arise from prior studies focusing on global measures of social anxiety symptoms that collapse across symptom domains. Fear of Negative Evaluation (FNE)—which refers to apprehension and distress arising from concerns about being judged negatively by others—is thought to reflect a core, defining symptom of social anxiety (Brown & Larson, 2009; Crone & Dahl, 2012; Dahl, 2004; Gerada, 2020; Nelemans et al., 2019; Parker et al., 2006; Steinberg & Morris, 2001; Westenberg et al., 2007), which may more closely relate to ERN variation in social settings. However, the broader cluster of symptoms associated with social anxiety also includes: social inhibition/avoidance, feelings of distress within social situations, and less confidence regarding social relationships (Hofmann, Heinrichs, & Moscovitch, 2004; Spokas & Cardaciotto, 2014; Yu, Zhou, Wang, & Tang, 2020), as well as potential overlap in symptom presentations from generalized anxiety disorder (Showraki et al., 2020). Thus, if the link between the ERN and social anxiety is primarily driven by FNE, then composite measures that collapse across FNE and other symptom domains could obscure such a link, leading to inconsistent findings across studies/samples.

Drawing on cognitive models of social anxiety and prior work investigating the ERN, we propose that ERN variation in social settings is more closely associated with FNE, as opposed to social anxiety symptoms more generally. Cognitive models propose a “vicious cycle” in which FNE leads to greater self-focus, self-monitoring and attentional biases toward social threats (i.e., errors/mistakes) within social settings, further exacerbating FNE as the result of a confirmation bias (Clark & Wells, 1995; Rapee & Heimberg, 1997). Thus, cognitive models explicitly predict a close association between FNE and self-monitoring for errors in social settings. Given that the ERN is an established index of self-monitoring for errors, we similarly hypothesize a close association between ERN magnitude within social settings and FNE (as opposed to social anxiety symptoms more generally). Indeed, neural responses associated with anticipated social feedback are correlated with FNE (Topel et al., 2021; Van der Molen et al., 2014), although prior work has not directly assessed the link between FNE and the ERN in social settings.

Adolescence is marked by increased social motivation, increased salience of peer relations (Brown & Larson, 2009), and heightened concern over peer evaluation (Parker et al., 2006; Steinberg & Morris, 2001). FNE not only reflects a hallmark symptom of social anxiety in cognitive models, but also exhibits a normative peak during adolescence (Brown & Larson, 2009; Crone & Dahl, 2012; Dahl, 2004; Gerada, 2020; Nelemans et al., 2019; Parker et al., 2006; Steinberg & Morris, 2001; Westenberg et al., 2007). Thus, our focus here on FNE, reflects a theoretically-relevant and developmentally-salient domain of social anxiety in adolescents.

The primary goal of the current study was to extend neurocognitive understandings of social anxiety in adolescence, as the first study to examine relations among the FNE subdomain of social anxiety symptoms and the ERN in peer presence. In the current study, a sample of adolescents (mean age: 16.98 years, SD = 0.45) residing in mainland China were assessed for social anxiety symptoms. EEG was recorded while performing a modified flanker task, twice, once while observed by a peer, and once while alone. Our hypotheses were twofold: 1) Given normative increases in peer importance during adolescence, participants would exhibit an overall increase in ERN magnitude in peer presence; 2) Higher FNE would predict ERN magnitudes in peer presence, whereas global social anxiety symptoms would exhibit diminished or non-significant associations with ERN in peer presence.

## Method

### 2.1. Participants and procedure

All participants in this study were recruited from grade 10 at Nanzheng High School, Hanzhong, mainland China. Exclusion criteria were 1) uncorrected visual impairments, 2) a history of head injury with loss of consciousness, and 3) neurological and developmental disorders. All parents and their adolescent children provided written informed consent/assent, and the research procedures were approved by the Shaanxi Normal University Human Research Ethics Board.

The current sample consisted of 30 adolescents (14 male, 16 female; mean age = 16.98 years, SD = 0.45, range: 15.68 – 17.91). One participant was excluded as the result of feeling unwell on the day of the assessment; two participants due to technical errors in data collection. In the final analyses, 27 participants (13 male, 14 female) were included in behavioral analyses, and 23 participants (10 male, 13 female) were included in EEG analyses.

Families that agreed to participate were invited to the lab. Upon arrival at the lab site, parents and their adolescent children were provided parental consent and child assent. Then parents were brought to another room to complete a demographic survey and the adolescents were asked to complete a modified flanker task (Eriksen & Eriksen, 1974) twice, once while they were being observed by a gender-matched peer who was actually the confederate (the social condition), and once while not being observed (the nonsocial condition). The order of the social and nonsocial conditions was counterbalanced across participants.

In the social condition, the child was informed that there was another participant who would conduct the experiment next to them and wanted to observe their performance to learn more about the task. The ages of the female and male confederates were 16.7 and 18.3 years old, respectively. During the social condition, the confederate was brought to the experiment room and introduced to the participant. To increase the ecological validity of the manipulation of social context and create a typical anxiety-provoking social situation, both confederates were trained to greet participants and make eye contact, but not to initiate any further oral communications, and only provide the simplest answers (i.e., yes, no, etc.) while being asked questions by participants. At the end of each block, the confederate read aloud the text feedback presented on the screen without any further comment.

In both conditions, steps were taken to minimize possible social observation effects of the experimenter being in the room while participants completed the flanker task, consistent with prior work (Barker et al., 2015; Voegler et al., 2018). The experimenter informed the participant that they would not observe their performance and was only there in case of unexpected technical problems. The experimenter then sat four meters away from the participant and pretended to read a book while the participant completed the flanker task.

### 2.2 Task, EEG Acquisition, and Preprocessing

#### Flanker Task

A modified version of the flanker task (Eriksen & Eriksen, 1974) was administered using e-prime software (Psychology Software Tools, Inc., Sharpsburg, PA). There were 8 blocks of 40 trials with a total of 320 trials. The target stimulus was a central arrowhead that pointed to the left or right and was flanked by four additional arrowheads (two on either side of the central arrowhead). Within each block, 50% of trials (20) consisted of target/flanker arrowheads pointing in the same direction (congruent) and 50% trials (20) consisted of arrowheads pointing in the opposite direction (incongruent); congruent and incongruent trials were presented in random order within each block. Arrowheads were presented in Lucida Console, size 240 font. All stimuli were presented for 200 ms followed by a 500 ms response window which ended upon the response. The response–stimulus interval jittered between 1000 to 1300 ms following the response or after 700 ms from stimulus onset. Figure 1 shows examples of stimulus sequences.

**Fig. 1.**
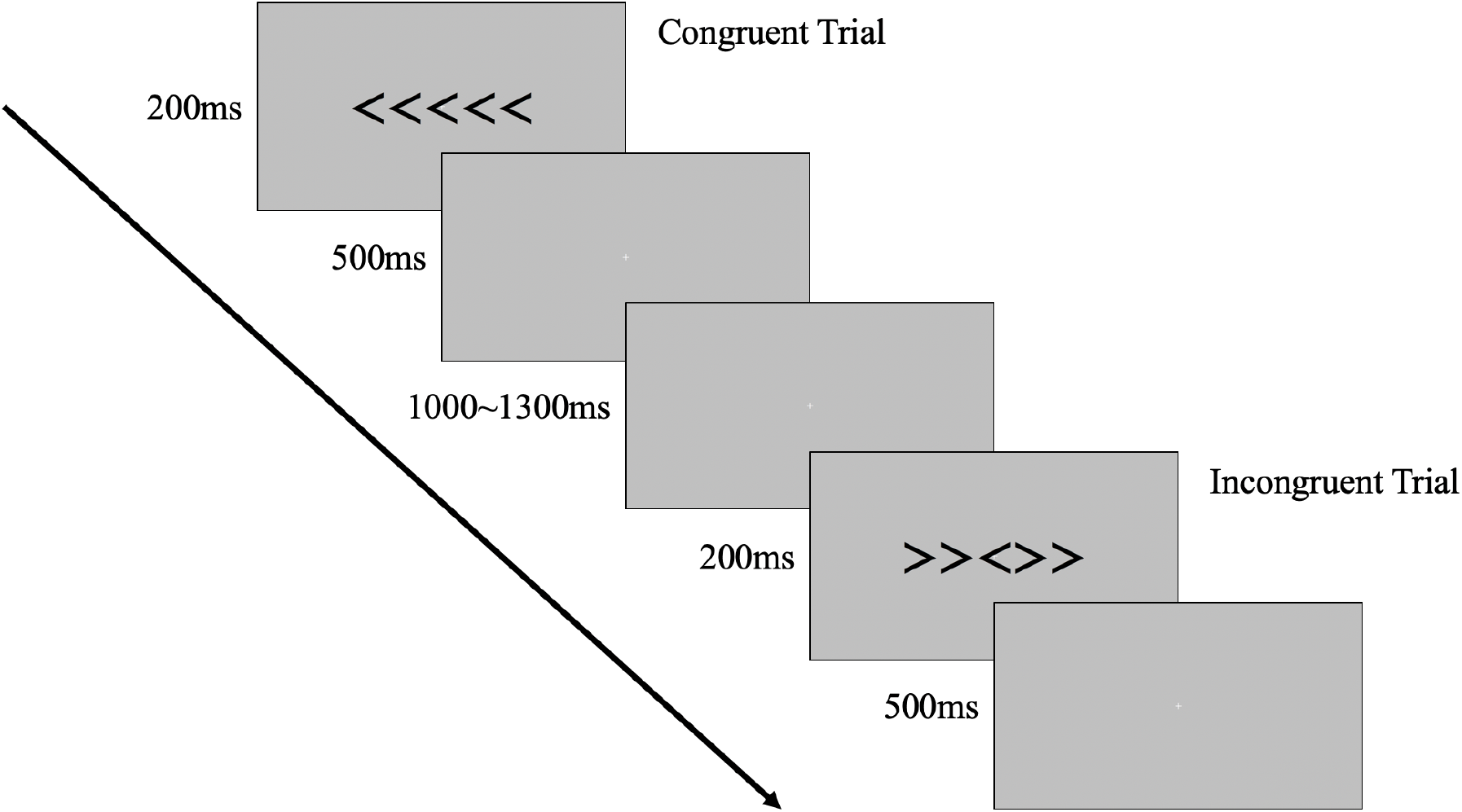
Schematic outline of Flanker task paradigm.

To ensure an adequate number of error trials for response-locked EEG analyses, block-level text-based feedback was provided at the end of each trial block contingent on the participant’s accuracy performance (Gehring, Liu, Orr, & Carp, 2012). If the accuracy for that block was below 75%, a message of “Please improve your accuracy!” was presented; If the accuracy was above 90%, a message of “Please respond a bit faster” was presented; if the accuracy was in-between 75% and 90%, a message of “Good job!” was presented. Block-level feedback was presented onscreen the same way within both the nonsocial and social conditions, however, the feedback text was additionally read aloud by the confederate in the social condition.

#### EEG Acquisition

Brain vision Recorder was used to record 64-channel EEG data, sampled at 500 Hz and acquired using a BrainVision ActiCHamp Amplifier, a high-impedance EEG system designed to maintain high signal-to-noise ratio at relatively high impedance levels. Before data collection, impedances were reduced to below 30 kΩ, consistent with manufacturer recommendations of 25-60 kΩ (Brain Products GmbH, 2016) and prior work (Laszlo et al., 2014). Data were online-referenced to the Cz electrode (note that data were rereferenced to the average of all electrodes as part of later preprocessing to recover EEG variation at/near Cz; Luck, 2014).

#### EEG Preprocessing

EEG data were preprocessed in MATLAB (Mathworks, Natick, MA) using the Maryland Analysis of Developmental EEG (MADE) Pipeline (Debnath et al., 2020). A detailed description of the preprocessing steps employed can be found in the original MADE publication (Debnath et al., 2020). Briefly, MADE consists of a set of MATLAB scripts and EEGLAB (Delorme & Makeig, 2004) functions/plugins, including FASTER (Nolan, Whelan, & Reilly, 2010) and Adjusted-ADJUST (Leach et al., 2020); MADE performs standard EEG preprocessing steps, including artifact removal via Independent component analysis (ICA) for artifact removal (Winkler, Debener, Müller, & Tangermann, 2015).

Following preprocessing via MADE, EEG data were epoched into 3000 ms segments starting 1000 ms before the response and baseline-corrected to the -400 to -200 ms window. Epochs that exceeded voltages ±100 μV, if the epochs were recorded from ocular channels (Fp1, Fp2, AF7, and AF8), they were further removed, and if the epochs were recorded from non-ocular channels, they were interpolated at the epoch level. All removed channels were interpolated using the spherical spline interpolation and then the data were referenced to the average of all electrodes. Epochs associated with anticipatory responses (RTs < 200 ms) were removed. Following preprocessing, two participants were removed from all further EEG analyses, as the result of having no usable data in one or more conditions.

#### Assessment of ERP Reliability

To set a minimum threshold for the number of trials required for inclusion in further analyses, ERN reliability in the current dataset was quantified— as a function of trial counts—using the ERP Reliability Analysis (ERA) toolbox (Clayson & Miller, 2017). These analyses demonstrated that a minimum cut-off of 7 error trials were needed to achieve a minimum internal consistency of ≥ 0.6. Applying this trial count threshold, two additional participants were removed from all further EEG analyses, as the result of having < 7 trials in one or more conditions. Altogether, 23 participants were included in the final EEG analyses. The mean numbers of trials-by-condition included in subsequent EEG analyses were as follows: 107.83 nonsocial-correct (SD = 22.07; range = 43–148), 28.52 nonsocial-error (SD = 11.67; range=7–49), 108.65 social-correct (SD = 22.21; range =58–146), and 29.83 social-error (SD = 12.54; range = 8–55).

### 2.3. Measures

#### Global Social Anxiety Symptom Severity

Self-reports of global social anxiety symptoms were assessed via the Social Phobia subscale of a Chinese translation of Screen for Child Anxiety Related Emotional Disorders (SCARED) (Birmaher et al., 1999; Su, Wang, Fan, Su, & Gao, 2008). Note that use of the term “Social Phobia” is consistent with the original manuscript introducing/validating the SCARED, designed to assesses anxiety symptoms in line with the Diagnostic and Statistical Manual (DSM) IV (American Psychiatric Association, 1994; Birmaher et al., 1999). The Social Phobia subscale consists of seven items (e.g., “It is hard for me to talk with people I don’t know well”) assessing global social anxiety symptom severity.

Each item is rated on a 3-point Likert scale, ranging from 0 (not true) to 2 (very true). Items are summed so that higher scores indicate higher levels of social anxiety symptom severity (total possible range: 0-14; scores ≥ 8 reflect the established clinical cut-off for social phobia [social anxiety] disorder (Birmaher et al., 1999). Prior work on the SCARED Social Phobia subscale demonstrates moderate to high internal consistency and test-retest reliability, and good discriminant validity within a Chinese population (Su et al., 2008). Normative data collected on the Social Phobia subscale, within a large sample of Chinese adolescents, are as follows: M = 4.10, *SD* = 3.22 (Su et al., 2008). The internal consistency of the Social Phobia subscale in the current sample was α = 0.85.

#### Fear of Negative Evaluation by Peers

Self-reports of Fear of Negative Evaluation (FNE) were assessed via the FNE subscale of a Chinese translation of the Social Anxiety Scale for Adolescents (SAS–A) (A. La Greca, 1998). The FNE subscale consists of eight items (e.g., “I worry about what other kids think of me”) assessing adolescents’ subjective experiences of fears, concerns, or worries regarding peers’ negative evaluations. Each item is rated on a 5-point Likert scale, ranging from 1 (not at all) to 5 (all the time). Items are summed so that higher scores indicate higher levels of FNE; (total possible range: 8-40). The Chinese version of the SAS has demonstrated appropriate internal consistency, test-retest reliability, and construct validity (Zhou, Xu, Inglés, Hidalgo, & La Greca, 2008). Normative data collected on the FNE subscale, within a large sample of Chinese adolescents, are as follows: M = 22.73, SD = 5.9 (Zhou et al., 2008). The internal consistency of the SAS–FNE subscale in the current sample was α = 0.94.

#### Flanker Task Behavior

Mean accuracy (ACC) and response time (RT) in each condition (nonsocial and social conditions) were calculated. Participants were required to have greater than 70% accuracy to be included in further behavioral and EEG analyses (all participants met this criterion, mean ACC = 87.23% for the nonsocial condition, and mean ACC = 87.04% for the social condition). Trial-level accuracy and RT data were extracted for further statistical analyses; RT data was first log-transformed to address positive skew (Luce, 1986).

#### ERPs

Only incongruent trials were included and analyzed (incongruent error trials for ERN, and incongruent correct trials for CRN) in order to isolate error-related effects and avoid stimulus-related confounds (errors are more common on incongruent trials in the flanker task). At the frontocentral electrode (FCz), where error-related brain activity is maximally negative (Meyer, Mehra, & Hajcak, 2021), trial-level mean amplitudes for the CRN and ERN were extracted within the 0 to 100 ms window following correct and incorrect responses, respectively.

### 2.4. Analytic Plan

For all behavioral and EEG analyses, we leveraged a mixed effects framework in R (R Core Team, 2013), given that this approach is capable of providing accurate and unbiased estimates, even in the presence low or unequal numbers of trials (Heise et al., 2022). For prediction of continuous outcome variables (ERPs, RT), Linear Mixed Effects (LME) modeling was employed, using the nlme package (Pinheiro et al., 2022); for prediction of binary accuracy data, a Generalized Linear Mixed Effects (GLME) modeling employed, using the glmer function in the lme4 package (Bates et al., 2014). In all analyses, categorical predictors were contrast coded (−1, +1), continuous predictors converted to Z scores, and effects estimated via maximum likelihood. To report p values, the anova function was used; interaction effects were proved via the emmeans package (Lenth, 2022).

#### Behavioral Analyses

Although not a primary focus of the current study, we carried out basic analyses of behavioral data (RT, Accuracy) for completeness. Separate mixed effects models were fit for accuracy and RT as dependent variables. For accuracy, a GLME model was estimated, with trial-level accuracy data as a binary outcome variable, effects of condition (nonsocial vs. social), congruency (congruent vs. incongruent), and their interaction were estimated as fixed effects, and participant entered as a random effect. For RT, a LME model was estimated, with RT as a continuous dependent variable, effects of condition (nonsocial vs. social), congruency (congruent vs. incongruent), response (correct vs. error), and their interactions estimated as fixed effects, and participants entered as a random effect.

#### ERP Analyses

To investigate effects of peer presence on ERN amplitude, we fit an LME model with trial-level ERP amplitudes as the outcome variable, effects of condition (nonsocial vs. social), response (correct vs. error), and their interaction estimated as fixed effects, and participant entered as a random effect. As described in the results, this analysis revealed a significant response-by-condition interaction, such that the ERP amplitudes on error trials (i.e., the ERN) but not correct trials (CRN) were increased in the social vs. nonsocial context. Thus, subsequent modelling focused on error trials only (ERN).

To examine effects of FNE vs. global anxiety symptoms on the ERN in peer presence/absence, we fit two separate LME models employing either FNE or global anxiety symptoms as predictors and directly compared relative model fit via Akaike information criterion (AIC) model selection. In each case, an LME model was fit, with trial-level ERN amplitudes as the outcome variable, condition (nonsocial vs. social) included as a fixed effect, and participant entered as a random effect. The FNE model additionally included FNE, as well its interaction with condition, as fixed effects; the global social anxiety symptoms model included global social anxiety symptoms, as well its interaction with condition, as fixed effects.

Given that the FNE model exhibited superior fit (see Results), we analyzed/interpreted results of this model; results of the global social anxiety model are reported in the supplement, for completeness. We additionally carried out a supplemental analysis in which global social anxiety symptoms were added to the FNE model as a control variable.

## Results

### 3.1. Self-report Questionnaires

The mean SCARED social anxiety score was 7.52 (*SD* = 3.50, range = 1 – 14), and the mean FNE score was 21.11 (*SD* = 8.74, range = 8 – 40). Although our analyses focus on continuous measures of social anxiety symptoms, consistent with a dimensional approach, it is worth noting that 12 of the 23 participants included in EEG analyses exceeded the established clinical cut-off for social phobia (social anxiety) disorder on the SCARED-SP (Birmaher et al., 1999). FNE was significantly correlated with SCARED social anxiety (*r* = .592, *p* < .001).

### 3.2. Behavioral Results

The mean (± standard deviation [SD]) accuracies in the nonsocial and social conditions were 87.23 ± 4.34% and 87.04 ± 4.61%, respectively (see Table 1). GLME modeling revealed a main effect of congruency, with participants responding less accurately in incongruent trials (*t*(16276) = 28.805, *p* < 0.001). There was no significant main effect of social condition (*t*(16276) = 0.650, *p* = 0.516), nor a congruency-by-condition interaction (*t*(16276) = 1.716, *p* = 0.086).

**Table 1.** Descriptive statistics characterizing behavior and ERPs as a function of condition

|  | Non-social condition | Social condition |
| --- | --- | --- |
| **Behavior Measures** | (N=27) | (N=27) |
| Accuracy (%) | 87.23 (4.34) | 87.04 (4.61) |
| Correct Reaction Time (ms) | 350.02 (35.78) | 350.20 (34.08) |
| Error Reaction time (ms) | 295.71 (30.50) | 294.31 (27.98) |
| **ERPs (μV)** | (N=23) | (N=23) |
| CRN | -0.87 (2.59) | -0.79 (2.48) |
| ERN | -4.52 (3.08) | -5.39 (3.44) |

Note. Values reflect mean scores (standard deviation in parentheses).

Mean RTs for the nonsocial and social conditions were 343.15 ± 36.36 ms and 343.02 ± 34.49 ms, respectively. LME modeling identified a main effect of response, with participants responding faster in error trials (*t*(16272) = 37.374, *p* < 0.001), and a main effect of congruency, with participants responding faster in congruent trials (*t*(16272) = -18.249, *p* < 0.001). Moreover, there was a response-by-congruency interaction effect (*t*(16272) = - 4.295, *p* < 0.001); within correct trials, participants responded slower in incongruent responses (*t*(16272) = 39.365, *p* < .001), within error trials, participants responded slower in incongruent responses (*t*(16272) = 7.273, *p* < .001). No significant main effect of social condition or other interaction effects were identified (all *p*s > .116). No correlations between behavioral data (accuracy and RT) and self-report questionnaires were statistically significant (all *p*s > .223).

### 3.3. ERP Results

Descriptive statistics for CRN and ERN in the nonsocial and social conditions are reported in Table 1. Figure 2 displays the ERP waveforms and scalp topography for the nonsocial and social conditions across participants at the FCz electrode (where the ERN was maximal).

**Fig. 2.**
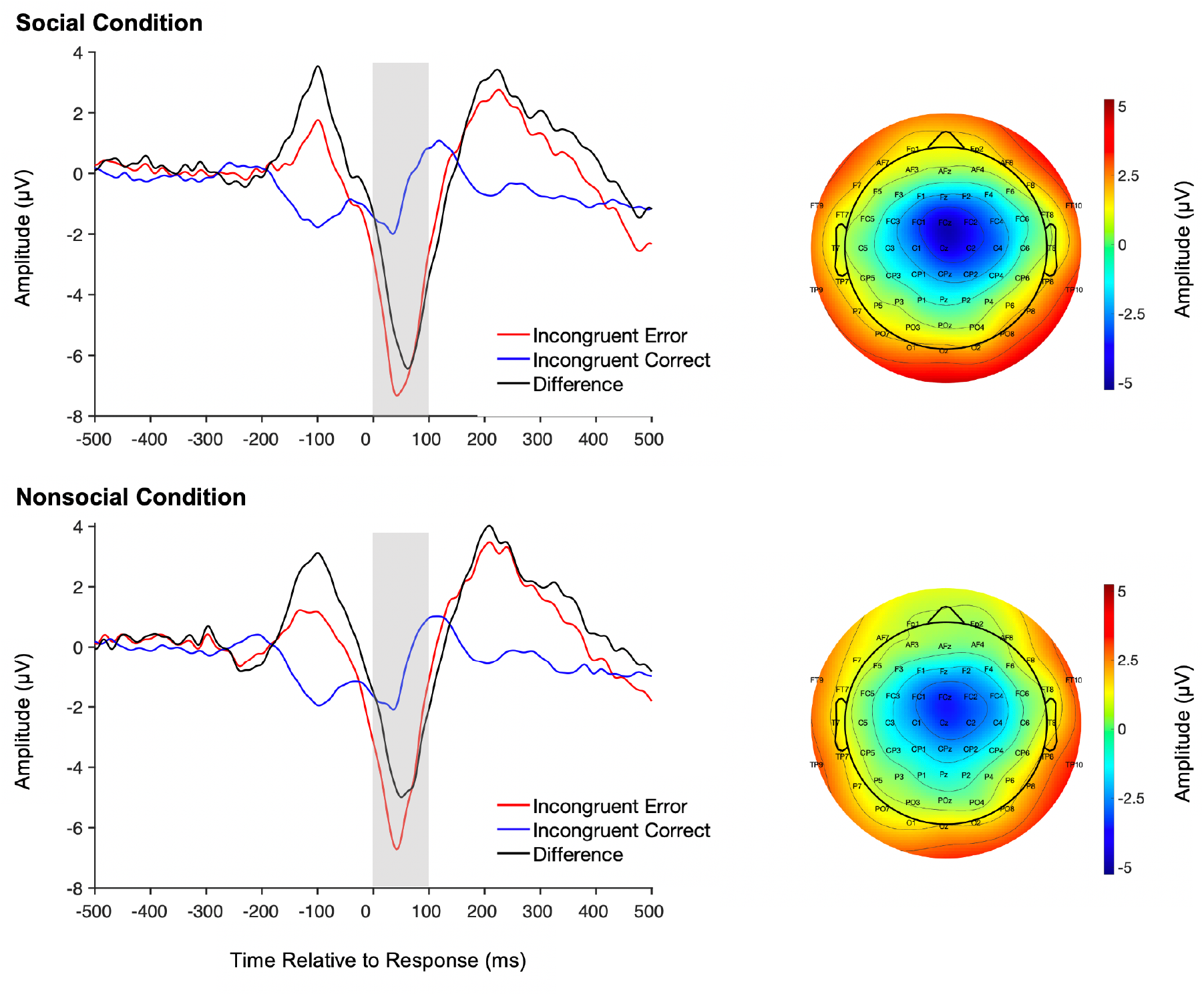
Averaged ERP and topographic plots for the social and nonsocial conditions. ERP plots depict averaged ERPs for incongruent-error trials (ERN: red), incongruent-correct trials (CRN: blue), and their difference (black), as a function of condition (top: social; bottom: nonsocial). Topographic plots corresponding to the shaded region within each ERP plot (0 – 100 ms) depict mean amplitude differences between incongruent-error and incongruent-correct trials, separately for each condition (top: social; bottom: nonsocial).

**Fig. 3.**
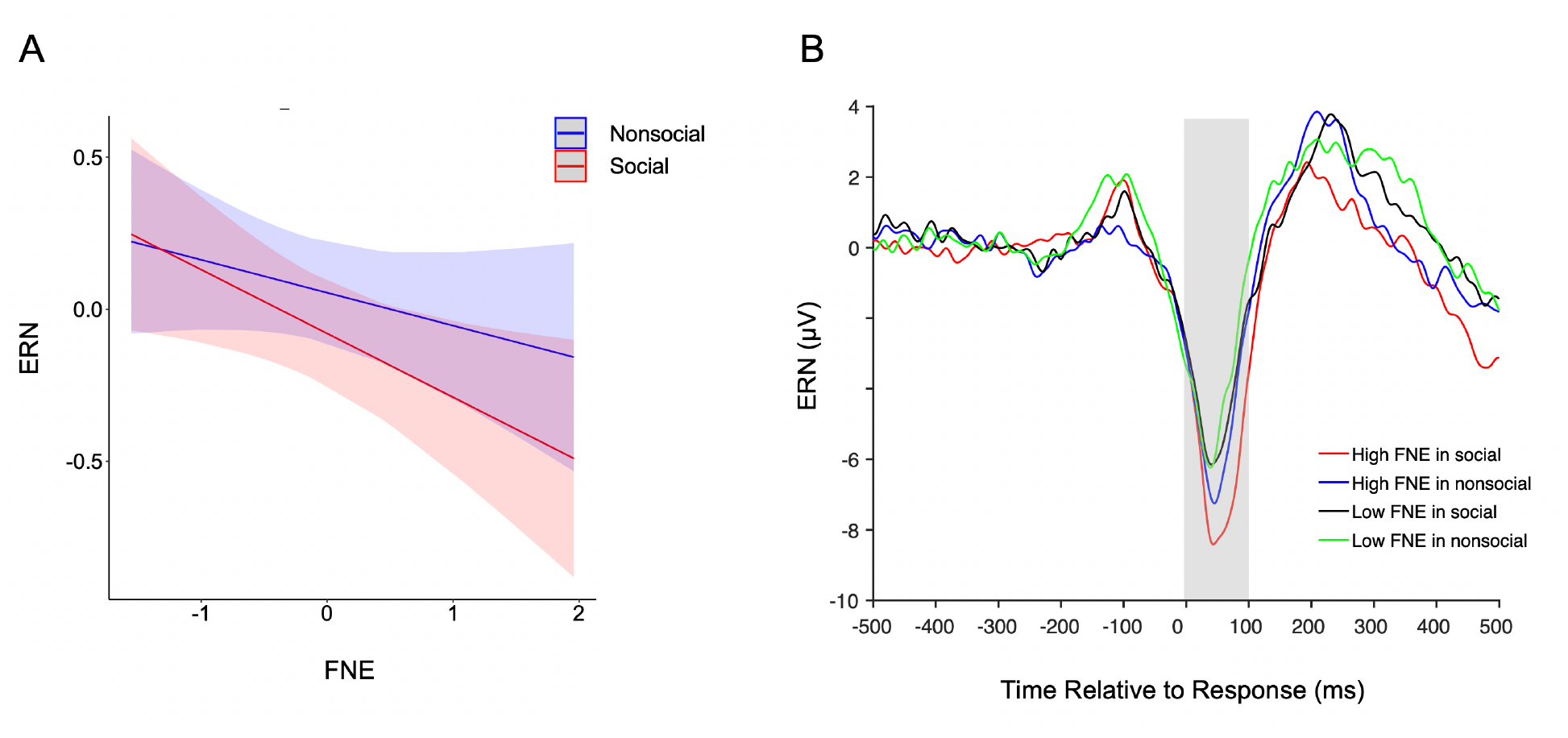
Association between FNE and ERN. A) Line plot depicting associations between FNE and the ERN, separately for the nonsocial (red) and social (blue) conditions. B) ERN ERP plots as a function of FNE group and condition (nonsocial vs. social). For plotting purposes only, participants were grouped into high and low FNE groups based on the median value of FNE scores (median = 18). ERN plots depict ERN ERPs for: the high FNE group in the social condition (red), high FNE group in the nonsocial condition (blue), low FNE group in the social condition (black), and low FNE group in the nonsocial condition (green). The shaded region reflects the 0 – 100 ms time window.

Results of the LME model assessing fixed effects of condition (nonsocial vs. social), response (correct vs. error), and their interaction, on trial-level ERP (ERN/CRN) amplitudes are reported in Table 2. This model revealed a significant main effect of response (*β* = 0.288, 95%CI [0.260, 0.317], *SE* = 0.014, *p* < .001), qualified by a response-by-condition interaction (*β* = 0.037, 95%CI [0.009, 0.065], *SE* = 0.014, *p* = .009); the nature of this interaction was such that the ERN was significantly larger in the social (compared to nonsocial) condition (*t*(6295) = 2.393, *p* = 0.017), whereas the CRN did not differ across conditions (*t*(6295) = -1.073, *p* = 0.283). Given that social context only exhibited a significant effect for error trials (ERN), and not correct trials (CRN), subsequent modelling focused only on error trials (ERN).

**Table 2.** **Model 1:** ERP ∼ Conditions * Response.

| Predictors | $\beta$ [95%CI] | SE | t | p |
| --- | --- | --- | --- | --- |
| (Intercept) | -0.167 [-0.301, -0.033] | 0.068 | -2.442 | .015* |
| Conditions | -0.023 [-0.051, 0.005] | 0.014 | -1.626 | .104 |
| Responses | 0.288 [0.260, 0.317] | 0.014 | 20.130 | < .001*** |
| Conditions*Responses | 0.037 [0.009, 0.065] | 0.014 | 2.620 | .009** |

Observations = 6321, N = 23.
Note: \* $p < .05$ , \*\* $p < .01$ , \*\*\* $p < .001$ .

We carried out Akaike information criterion (AIC) model selection to compare the following LME models: 1) the FNE model, defined by fixed effects of FNE, condition (nonsocial vs. social), and their interaction, on trial-level ERN amplitudes; 2) the global social anxiety model, defined by fixed effects of global social anxiety, condition (nonsocial vs. social), and their interaction, on trial-level ERN amplitudes. AIC model selection revealed that FNE model, carrying 81% of the cumulative model weight, had lower AIC (AIC = 8933.95) than global social anxiety model (AIC = 8936.83), indicating that the FNE model provided a better fit to the data. Given that the FNE model exhibited superior fit, subsequent analyses focused on interpretation of the results of this model (however, for completeness, see the supplement for results of the global social anxiety model).

Results of the LME model assessing fixed effects of condition (nonsocial vs. social), FNE, and their interaction, on trial-level ERN amplitudes are reported in Table 3. This model revealed a significant main effect of condition (*t*(1317) = -2.672, *p* = 0.008), qualified by a condition-by-FNE interaction (*β* = -0.052, 95%CI [-0.102, -0.002], *SE* = 0.026, *p* = .042); the nature of this interaction was such that higher levels of FNE were more strongly associated with a larger (more negative) ERN in the social (compared to nonsocial) condition (*t*(1317) = 2.032, *p*= 0.042). We additionally carried out a supplemental analysis in which global social anxiety symptoms were added to the FNE model, described above, as a control variable. As reported in the supplement, the condition-by-FNE interaction remained significant when controlling for global social anxiety symptoms (*β* = -0.052, 95%CI [-0.102, -0.002], *SE* = 0.026, *p* = .042), STATS, demonstrating that FNE predicted unique variance in ERN, while controlling for global social anxiety symptoms.

**Table 3.** **Model 2:** ERN ∼ Conditions * FNE

| Predictors | $\beta$ [95%CI] | SE | t | p |
| --- | --- | --- | --- | --- |
| (Intercept) | -0.461 [-0.623, -0.299] | 0.083 | -5.580 | < .001*** |
| Conditions | -0.069 [-0.119, -0.018] | 0.026 | -2.672 | .008** |
| FNE | -0.163 [-0.338, 0.012] | 0.084 | -1.935 | .067 |
| Conditions*FNE | -0.052 [-0.102, -0.002] | 0.026 | -2.032 | .042* |

Observations = 1342, N = 23. FNE = Fear of Negative Evaluation.
Note: \* $p < .05$ , \*\* $p < .01$ , \*\*\* $p < .001$ .

## Discussion

Consistent with our hypotheses, we observed an overall increase in ERN magnitude in peer presence. Comparing the statistical models involving either FNE or global social anxiety symptoms to predict peer/alone ERN variation revealed a better fit for the FNE model. Moreover, higher levels of FNE were predictive of an increased ERN in the peer (vs. alone) setting— an effect that held even when controlling for global social anxiety symptoms. In contrast, global social anxiety symptoms did not significantly relate to the ERN. Results of the current study are consistent with cognitive models of social anxiety (Clark & Wells, 1995; Rapee & Heimberg, 1997), which propose a close link between FNE and self-monitoring for errors within social settings, providing a crucial neurocognitive extension of prior work. Moreover, these findings are consistent with developmental theory describing an increase in the importance of peers and normatively high levels of FNE during adolescence (Brown & Larson, 2009; Crone & Dahl, 2012; Dahl, 2004; Gerada, 2020; Nelemans et al., 2019; Parker et al., 2006; Steinberg & Morris, 2001; Westenberg et al., 2007). These findings have implications for theory and measurement of adolescent social anxiety, including the importance of considering error sensitivity (Chong & Meyer, 2019) and the possibility of targeting error-related processing (Meyer et al., 2020) in adolescent social anxiety. These data also highlight the ERN’s utility as a neurocognitive index of self-monitoring within social settings, which complements self-report measures and could be leveraged by future studies seeking to examine causal mechanisms predicted by cognitive models of social anxiety. We discuss these ideas in further detail below.

Given that social anxiety is characterized by a cluster of heterogeneous symptoms (Inderbitzen-Nolan & Walters, 2000; A. M. La Greca & Lopez, 1998; Watson & Friend, 1969), it is reasonable to assume that no single neurocognitive measure would relate to all social anxiety symptoms equally (or at least not reliably across study samples); results of the current study are in line with this notion. Moreover, our data suggest that successful identification of links between symptom domains and neurocognitive measures requires precision at the level of symptom assessment as well as taking into account the situational context of neural assessment and the broader developmental context. Toward these ends, the current study focused on relations between a developmentally-salient symptom domain of social anxiety—FNE (e.g., A. M. La Greca & Lopez, 1998; Watson & Friend, 1969) and neural activity associated with error-monitoring (self-monitoring) under peer observation (e.g., Hajcak et al., 2005; Kim et al., 2005), within the context of adolescence—when FNE and the importance of peers reach peak levels (Brown & Larson, 2009; Crone & Dahl, 2012; Dahl, 2004; Gerada, 2020; Nelemans et al., 2019; Parker et al., 2006; Steinberg & Morris, 2001; Westenberg et al., 2007).

Importantly, the current study demonstrates that an increased ERN in social settings is associated with the FNE symptom domain in adolescents. Yet, we do not suggest that the ERN reflects the neurocognitive basis of FNE per se. Prior work demonstrates that the ERN indexes a neurocognitive process underlying error-monitoring (self-monitoring) (Agam et al., 2011; Buzzell et al., 2017a; Dehaene et al., 1994; Falkenstein et al., 1991; Gehring et al., 1993; Miltner et al., 1997; Ullsperger, Fischer, Nigbur, & Endrass, 2014). Similarly, we suggest that an increased ERN in social settings indexes increased self-monitoring. As noted, cognitive models propose a vicious cycle whereby greater FNE leads to greater self-monitoring within social settings, which in turn further strengthens/worsens FNE (Clark & Wells, 1995; Rapee & Heimberg, 1997). Thus, we argue that the link between FNE and the ERN in peer presence reflects a neurocognitive extension of cognitive models of social anxiety. These models further state that social anxiety is specifically associated with increased self-monitoring for negative aspects of one’s behavior (e.g., errors) as opposed to a general increase in self-monitoring (Clark & Wells, 1995; Rapee & Heimberg, 1997). Our data provide converging neural evidence to support this supposition, as peer presence and FNE predicted larger increases in the ERN (self-monitoring on error trials), as opposed to the CRN (self-monitoring on correct trials). As noted by cognitive models (Clark & Wells, 1995; Rapee & Heimberg, 1997), increased self-monitoring of negative aspects of one’s behavior could explain why social anxiety is maintained/worsened across repeated social encounters in daily life: social encounters are experienced through the lens of one’s (neuro-) cognitive biases. A bias towards greater self-monitoring for errors in social settings may serve to confirm one’s initial fears of negative evaluation, strengthening/worsening FNE.

This is the first study to directly investigate relations between FNE and the ERN in peer presence among adolescents. Given our cross-sectional study design, it remains unknown whether the association between FNE and ERN remains stable across development. Emerging developmental theory on the nature of anxiety-ERN relations argues that whereas the ERN-anxiety link in childhood is primarily driven by external threats/fears, ERN-anxiety relations fundamentally change with the transition to adolescence—to self-conscious shyness and worry about behavioral competence and social evaluation (Meyer, 2017, 2022). Meyer et al. have shown that adolescent ERN-anxiety relations are primarily driven by social anxiety symptoms, as opposed to symptoms of other anxiety disorders (Meyer et al., 2018). Moreover, normative ERN increases during the transition to adolescence are mediated by (normative) increases in subclinical social anxiety symptoms (Meyer et al., 2018). Based on the results of the current study, taken together with the normative increase in FNE during the transition to adolescence (Gerada, 2020; Nelemans et al., 2019; Westenberg et al., 2007), we propose that ERN variation becomes more closely coupled with individual differences in FNE across adolescence. Longitudinal studies that measure ERN and FNE across the peri-adolescent window are needed to evaluate this proposal.

The current study has several implications for clinical neuroscience research and efforts to develop novel assessment approaches. Broadly, the current findings provide additional emphasis on the importance of understanding error sensitivity (Chong & Meyer, 2019) and specifically targeting error-related processing (Meyer et al., 2020) in pediatric anxiety, particularly during adolescence. Moreover, assessment of the ERN in peer presence may reflect a promising neural marker, indicating the degree to which an individual exhibits greater self-monitoring in social settings. As such the ERN could provide a complement to self-report measures and be particularly useful in settings or participant groups more likely to be impacted by metacognitive awareness, social desirability, or recall biases (Brewin et al., 1993; Van de Mortel, 2008). While we argue that the ERN underlies self-monitoring, as opposed to a direct index of FNE, cognitive models of social anxiety (Clark & Wells, 1995; Rapee & Heimberg, 1997) and converging neurocognitive evidence from the current study, establish a close association between self-monitoring in peer presence and FNE. Thus, the ERN could also be employed as an (indirect) neural marker of FNE levels. Going one step further, an intriguing possibility is that repeated assessments of FNE and ERN, at the intraindividual level, could provide the means to directly capture the “vicious cycle” proposed by cognitive models of social anxiety (Clark & Wells, 1995; Rapee & Heimberg, 1997). However, further work examining intraindividual relations among FNE and ERN are needed, to include dissociating trait-vs. state-based associations in these constructs. There is also a need for additional work studying these constructs via methods capable of identifying causal pathways (e.g., longitudinal designs or causal manipulation studies). Such work would provide strong evidence for whether/how FNE and the ERN causally influence one another, and a critical test of whether these constructs are implicated in a “vicious cycle” in which they mutually strengthen/worsen one another.

### Strengths, Limitations and Future Directions

Strengths of the current study include the targeted assessment of a theoretically relevant and developmentally-salient symptom domain of social anxiety—FNE, inclusion of a non-WEIRD (Western, Educated, Industrialized, Rich and Democratic) (Henrich, Heine, & Norenzayan, 2010) sample of adolescents, and at the theoretical level, providing a neurocognitive extension of cognitive models of social anxiety. Yet, this study has several limitations. The first limitation is that our sample size was relatively small, resulting in low statistical power to detect small effects; thus, results of the current report should be interpreted in light of the small sample size. We were also not sufficiently powered to test whether biological sex or gender moderates observed links between ERN, FNE and social context. Similarly, the age range of participants was selected to be narrow in the current study to minimize possible age-related effects, as we would similarly be underpowered to test whether age moderates links between ERN, FNE and social context. Therefore, future work is needed to replicate our findings within a larger sample, and should adopt a broader sampling strategy to investigate whether the reported results are moderated by biological sex, gender, or age. Note that additional limitations of the current study are that, although a priori, hypotheses were not pre-registered, and there were no planned analyses of power to determine sample size. Thus, attempts to replicate the current results could further benefit from pre-registering hypotheses, and leverage effect sizes reported in the current study to carry-out a priori power analyses. It is also worth noting that the current study employed a standard, lab-based assessment of error monitoring via the Flanker task, assessed at a single point in time. Future work could extend this approach via repeated, longitudinal assessments within the lab, or even work to develop recording/analysis protocols that would allow for more continuous, in-vivo assessment of these constructs outside the laboratory (Caricato et al., 2020). While a strength of the current study is the inclusion of a non-WEIRD sample, it remains unknown whether associations between social anxiety symptomology and the ERN in peer presence are stable across cultures; cross-cultural studies are needed. Lastly, when interpreting the findings, the cross-sectional design of the current study should be taken into account, since it does not allow us to draw conclusions on the causal direction of associations between FNE and ERN. Regardless, the current study has important implications for clinical theory and social anxiety research; it demonstrates the utility of mapping neurocognitive construct(s) onto specific symptom domain(s) of social anxiety, and highlights the importance of carefully considering precision in symptom assessment, the situational context of neural measurement, and the developmental backdrop in which such work is conducted.

## Acknowledgments

The authors would like to thank all families who participated in the study for their time and contribution.

## Ethical standards

The authors assert that all procedures contributing to this work comply with the ethical standards of the relevant national and institutional committees on human experimentation and with the Helsinki Declaration of 1975, as revised in 2008.

## References

Agam, Y., Hämäläinen, M. S., Lee, A. K., Dyckman, K. A., Friedman, J. S., Isom, M., Makris, N., & Manoach, D. S. (2011). Multimodal neuroimaging dissociates hemodynamic and electrophysiological correlates of error processing. Proceedings of the National Academy of Sciences, 108(42), 17556–17561.

Allain, S., Carbonnell, L., Falkenstein, M., Burle, B., & Vidal, F. (2004). The modulation of the Ne-like wave on correct responses foreshadows errors. Neuroscience Letters, 372(1–2), 161–166.

American Psychiatric Association. (1994). Diagnostic and Statistical Manual of Mental Disorders, 4th edn revised. Washington DC: American Psychiatric Association, 317–392.

Barker, T. V., Troller-Renfree, S., Pine, D. S., & Fox, N. A. (2015). Individual differences in social anxiety affect the salience of errors in social contexts. Cognitive, Affective, & Behavioral Neuroscience, 15(4), 723–735.

Bates, D., Mächler, M., Bolker, B., & Walker, S. (2014). Fitting linear mixed-effects models using lme4. ArXiv Preprint ArXiv:1406.5823.

Birmaher, B., Brent, D. A., Chiappetta, L., Bridge, J., Monga, S., & Baugher, M. (1999). Psychometric properties of the Screen for Child Anxiety Related Emotional Disorders (SCARED): A replication study. Journal of the American Academy of Child & Adolescent Psychiatry, 38(10), 1230–1236.

Brain Products GmbH. (2016). ActiCHamp Operating Instructions. www.brainproducts.com

Brewin, C. R., Andrews, B., & Gotlib, I. H. (1993). Psychopathology and early experience: A reappraisal of retrospective reports. Psychological Bulletin, 113(1), 82.

Brown, B. B., & Larson, J. (2009). Peer relationships in adolescence.

Buzzell, G. A., Richards, J. E., White, L. K., Barker, T. V., Pine, D. S., & Fox, N. A. (2017). Development of the error-monitoring system from ages 9–35: Unique insight provided by MRI-constrained source localization of EEG. Neuroimage, 157, 13–26.

Buzzell, G. A., Troller-Renfree, S. V., Barker, T. V., Bowman, L. C., Chronis-Tuscano, A., Henderson, H. A., Kagan, J., Pine, D. S., & Fox, N. A. (2017). A neurobehavioral mechanism linking behaviorally inhibited temperament and later adolescent social anxiety. Journal of the American Academy of Child & Adolescent Psychiatry, 56(12), 1097–1105.

Caricato, A., Della Marca, G., Ioannoni, E., Silva, S., Benzi Markushi, T., Stival, E., Biasucci, D. G., Montano, N., Gelormini, C., & Melchionda, I. (2020). Continuous EEG monitoring by a new simplified wireless headset in intensive care unit. BMC Anesthesiology, 20(1), 1–6.

Chong, L. J., & Meyer, A. (2019). Understanding the link between anxiety and a neural marker of anxiety (the error-related negativity) in 5 to 7 year-old children. Developmental Neuropsychology, 44(1), 71–87.

Clark, D. M., & Wells, A. (1995). A cognitive model. Social Phobia: Diagnosis, Assessment, and Treatment, 69, 1025.

Clayson, P. E., & Miller, G. A. (2017). ERP Reliability Analysis (ERA) Toolbox: An open-source toolbox for analyzing the reliability of event-related brain potentials. International Journal of Psychophysiology, 111, 68–79.

Crone, E. A., & Dahl, R. E. (2012). Understanding adolescence as a period of social–affective engagement and goal flexibility. Nature Reviews Neuroscience, 13(9), 636–650.

Dahl, R. E. (2004). Adolescent brain development: A period of vulnerabilities and opportunities. Keynote address. Annals of the New York Academy of Sciences, 1021(1), 1–22.

Debnath, R., Buzzell, G. A., Morales, S., Bowers, M. E., Leach, S. C., & Fox, N. A. (2020). The Maryland analysis of developmental EEG (MADE) pipeline. Psychophysiology, 57(6), e13580.

Dehaene, S., Posner, M. I., & Tucker, D. M. (1994). Localization of a neural system for error detection and compensation. Psychological Science, 5(5), 303–305.

Delorme, A., & Makeig, S. (2004). EEGLAB: an open source toolbox for analysis of single-trial EEG dynamics including independent component analysis. Journal of Neuroscience Methods, 134(1), 9–21.

Endrass, T., Riesel, A., Kathmann, N., & Buhlmann, U. (2014). Performance monitoring in obsessive–compulsive disorder and social anxiety disorder. Journal of Abnormal Psychology, 123(4), 705.

Eriksen, B. A., & Eriksen, C. W. (1974). Effects of noise letters upon the identification of a target letter in a nonsearch task. Perception & Psychophysics, 16(1), 143–149.

Falkenstein, M., Hohnsbein, J., Hoormann, J., & Blanke, L. (1991). Effects of crossmodal divided attention on late ERP components. II. Error processing in choice reaction tasks. Electroencephalography and Clinical Neurophysiology, 78(6), 447–455.

Ford, J. M. (1999). Schizophrenia: The broken P300 and beyond. Psychophysiology, 36(6), 667–682.

Gehring, W. J., Goss, B., Coles, M. G., Meyer, D. E., & Donchin, E. (1993). A neural system for error detection and compensation. Psychological Science, 4(6), 385–390.

Gehring, W. J., Liu, Y., Orr, J. M., & Carp, J. (2012). The error-related negativity (ERN/Ne).

Gerada, A. (2020). The Longitudinal Association Between Body Image Dissatisfaction, Social Anxiety, and Fear of Negative Evaluation in Adolescents [PhD Thesis]. Université d’Ottawa/University of Ottawa.

Hajcak, G., & Foti, D. (2008). Errors are aversive: Defensive motivation and the error-related negativity. Psychological Science, 19(2), 103–108.

Hajcak, G., Klawohn, J., & Meyer, A. (2019). The utility of event-related potentials in clinical psychology. Annual Review of Clinical Psychology, 15, 71–95.

Hajcak, G., McDonald, N., & Simons, R. F. (2003). To err is autonomic: Error-related brain potentials, ANS activity, and post-error compensatory behavior. Psychophysiology, 40(6), 895–903.

Hajcak, G., Moser, J. S., Yeung, N., & Simons, R. F. (2005). On the ERN and the significance of errors. Psychophysiology, 42(2), 151–160.

Heise, M. J., Mon, S. K., & Bowman, L. C. (2022). Utility of linear mixed effects models for event-related potential research with infants and children. Developmental Cognitive Neuroscience, 54, 101070.

Henrich, J., Heine, S. J., & Norenzayan, A. (2010). The weirdest people in the world? Behavioral and Brain Sciences, 33(2–3), 61–83.

Hofmann, S. G., Heinrichs, N., & Moscovitch, D. A. (2004). The nature and expression of social phobia: Toward a new classification. Clinical Psychology Review, 24(7), 769–797.

Inderbitzen-Nolan, H. M., & Walters, K. S. (2000). Social Anxiety Scale for Adolescents: Normative data and further evidence of construct validity. Journal of Clinical Child Psychology, 29(3), 360–371.

Kessler, R. C., Berglund, P., Demler, O., Jin, R., Merikangas, K. R., & Walters, E. E. (2005). Lifetime prevalence and age-of-onset distributions of DSM-IV disorders in the National Comorbidity Survey Replication. Archives of General Psychiatry, 62(6), 593–602.

Kim, E. Y., Iwaki, N., Uno, H., & Fujita, T. (2005). Error-related negativity in children: Effect of an observer. Developmental Neuropsychology, 28(3), 871–883.

Kujawa, A., Weinberg, A., Bunford, N., Fitzgerald, K. D., Hanna, G. L., Monk, C. S., Kennedy, A. E., Klumpp, H., Hajcak, G., & Phan, K. L. (2016). Error-related brain activity in youth and young adults before and after treatment for generalized or social anxiety disorder. Progress in Neuro-Psychopharmacology and Biological Psychiatry, 71, 162–168.

La Greca, A. (1998). Manual for the social anxiety scales for children and adolescents. Miami, FL: University of Miami.

La Greca, A. M., & Lopez, N. (1998). Social anxiety among adolescents: Linkages with peer relations and friendships. Journal of Abnormal Child Psychology, 26(2), 83–94.

Laszlo, S., Ruiz-Blondet, M., Khalifian, N., Chu, F., & Jin, Z. (2014). A direct comparison of active and passive amplification electrodes in the same amplifier system. Journal of Neuroscience Methods, 235, 298–307.

Leach, S. C., Morales, S., Bowers, M. E., Buzzell, G. A., Debnath, R., Beall, D., & Fox, N. A. (2020). Adjusting ADJUST: Optimizing the ADJUST algorithm for pediatric data using geodesic nets. Psychophysiology, 57(8). https://doi.org/10.1111/psyp.13566

Lenth, R. V. (2022). emmeans: Estimated Marginal Means, aka Least-Squares Means. https://CRAN.R-project.org/package=emmeans

Luce, R. D. (1986). Response times: Their role in inferring elementary mental organization. Oxford University Press on Demand.

Luck, S. J. (2014). An introduction to the event-related potential technique. MIT press.

Masaki, H., Maruo, Y., Meyer, A., & Hajcak, G. (2017). Neural correlates of choking under pressure: Athletes high in sports anxiety monitor errors more when performance is being evaluated. Developmental Neuropsychology, 42(2), 104–112.

Meyer, A. (2017). A biomarker of anxiety in children and adolescents: A review focusing on the error-related negativity (ERN) and anxiety across development. Developmental Cognitive Neuroscience, 27, 58–68. https://doi.org/10.1016/j.dcn.2017.08.001

Meyer, A. (2022). On the relationship between the error-related negativity and anxiety in children and adolescents: From a neural marker to a novel target for intervention. Psychophysiology, e14050.

Meyer, A., Carlton, C., Chong, L. J., & Wissemann, K. (2019). The presence of a controlling parent is related to an increase in the error-related negativity in 5–7 year-old children. Journal of Abnormal Child Psychology, 47(6), 935–945.

Meyer, A., Carlton, C., Crisler, S., & Kallen, A. (2018). The development of the error-related negativity in large sample of adolescent females: Associations with anxiety symptoms. Biological Psychology, 138, 96–103.

Meyer, A., Gibby, B., Wissemann, K., Klawohn, J., Hajcak, G., & Schmidt, N. B. (2020). A brief, computerized intervention targeting error sensitivity reduces the error-related negativity. Cognitive, Affective, & Behavioral Neuroscience, 20(1), 172–180.

Meyer, A., Mehra, L., & Hajcak, G. (2021). Error-related negativity predicts increases in anxiety in a sample of clinically anxious female children and adolescents over 2 years. Journal of Psychiatry and Neuroscience, 46(4), E472–E479.

Miltner, W. H., Braun, C. H., & Coles, M. G. (1997). Event-related brain potentials following incorrect feedback in a time-estimation task: Evidence for a “generic” neural system for error detection. Journal of Cognitive Neuroscience, 9(6), 788–798.

Morrison, A. S., & Heimberg, R. G. (2013). Social anxiety and social anxiety disorder. Annual Review of Clinical Psychology, 9, 249–274.

Moser, J., Moran, T., Schroder, H., Donnellan, B., & Yeung, N. (2013). On the relationship between anxiety and error monitoring: A meta-analysis and conceptual framework. Frontiers in Human Neuroscience, 7, 466.

Nelemans, S. A., Meeus, W. H., Branje, S. J., Van Leeuwen, K., Colpin, H., Verschueren, K., & Goossens, L. (2019). Social Anxiety Scale for Adolescents (SAS-A) Short Form: Longitudinal measurement invariance in two community samples of youth. Assessment, 26(2), 235–248.

Nolan, H., Whelan, R., & Reilly, R. B. (2010). FASTER: fully automated statistical thresholding for EEG artifact rejection. Journal of Neuroscience Methods, 192(1), 152–162.

Olvet, D. M., & Hajcak, G. (2008). The error-related negativity (ERN) and psychopathology: Toward an endophenotype. Clinical Psychology Review, 28(8), 1343–1354.

Parker, J. G., Rubin, K. H., Erath, S. A., Wojslawowicz, J. C., & Buskirk, A. A. (2006). Peer relationships, child development, and adjustment: A developmental psychopathology perspective.

Pineles, S. L., & Mineka, S. (2005). Attentional biases to internal and external sources of potential threat in social anxiety. Journal of Abnormal Psychology, 114(2), 314.

Pinheiro, J., Bates, D., & R Core Team. (2022). nlme: Linear and Nonlinear Mixed Effects Models. https://CRAN.R-project.org/package=nlme

Rapee, R. M., & Heimberg, R. G. (1997). A cognitive-behavioral model of anxiety in social phobia. Behaviour Research and Therapy, 35(8), 741–756.

Schillinger, F. L., De Smedt, B., & Grabner, R. H. (2016). When errors count: An EEG study on numerical error monitoring under performance pressure. Zdm, 48(3), 351–363.

Showraki, M., Showraki, T., & Brown, K. (2020). Generalized anxiety disorder: Revisited. Psychiatric Quarterly, 91(3), 905–914.

Spokas, M. E., & Cardaciotto, L. (2014). Heterogeneity within social anxiety disorder. In The Wiley Blackwell handbook of social anxiety disorder (pp. 247–267). Wiley Blackwell.

Steinberg, L., & Morris, A. S. (2001). Adolescent development. Annual Review of Psychology, 52(1), 83–110.

Su, L., Wang, K., Fan, F., Su, Y., & Gao, X. (2008). Reliability and validity of the screen for child anxiety related emotional disorders (SCARED) in Chinese children. Journal of Anxiety Disorders, 22(4), 612–621.

Suchan, B., Jokisch, D., Skotara, N., & Daum, I. (2007). Evaluation-related frontocentral negativity evoked by correct responses and errors. Behavioural Brain Research, 183(2), 206–212.

Team, R. C. & others. (2013). R: A language and environment for statistical computing.

Topel, S., van Noordt, S. J., Willner, C. J., Banz, B. C., Wu, J., Castagna, P., Kortink, E. D., Van der Molen, M. J., & Crowley, M. J. (2021). As they wait: Anticipatory neural response to evaluative peer feedback varies by pubertal status and social anxiety. Developmental Cognitive Neuroscience, 51, 101004.

Ullsperger, M., Fischer, A. G., Nigbur, R., & Endrass, T. (2014). Neural mechanisms and temporal dynamics of performance monitoring. Trends in Cognitive Sciences, 18(5), 259–267.

Van de Mortel, T. F. (2008). Faking it: Social desirability response bias in self-report research. Australian Journal of Advanced Nursing, The, 25(4), 40–48.

Van der Molen, M. J., Poppelaars, E. S., Van Hartingsveldt, C. T., Harrewijn, A., Gunther Moor, B., & Westenberg, P. M. (2014). Fear of negative evaluation modulates electrocortical and behavioral responses when anticipating social evaluative feedback. Frontiers in Human Neuroscience, 7, 936.

Van Meel, C. S., & Van Heijningen, C. A. (2010). The effect of interpersonal competition on monitoring internal and external error feedback. Psychophysiology, 47(2), 213–222.

Vidal, F., Burle, B., Bonnet, M., Grapperon, J., & Hasbroucq, T. (2003). Error negativity on correct trials: A reexamination of available data. Biological Psychology, 64(3), 265–282.

Vidal, F., Hasbroucq, T., Grapperon, J., & Bonnet, M. (2000). Is the ‘error negativity’specific to errors? Biological Psychology, 51(2–3), 109–128.

Voegler, R., Peterburs, J., Lemke, H., Ocklenburg, S., Liepelt, R., & Straube, T. (2018). Electrophysiological correlates of performance monitoring under social observation in patients with social anxiety disorder and healthy controls. Biological Psychology, 132, 71–80.

Watson, D., & Friend, R. (1969). Measurement of social-evaluative anxiety. Journal of Consulting and Clinical Psychology, 33(4), 448.

Westenberg, P. M., Gullone, E., Bokhorst, C. L., Heyne, D. A., & King, N. J. (2007). Social evaluation fear in childhood and adolescence: Normative developmental course and continuity of individual differences. British Journal of Developmental Psychology, 25(3), 471–483.

Winkler, I., Debener, S., Müller, K.-R., & Tangermann, M. (2015). On the influence of high-pass filtering on ICA-based artifact reduction in EEG-ERP. 2015 37th Annual International Conference of the IEEE Engineering in Medicine and Biology Society (EMBC), 4101–4105.

Yu, M., Zhou, H., Wang, M., & Tang, X. (2020). The heterogeneity of social anxiety symptoms among Chinese adolescents: Results of latent profile analysis. Journal of Affective Disorders, 274, 935–942.

Zhou, X., Xu, Q., Inglés, C. J., Hidalgo, M. D., & La Greca, A. M. (2008). Reliability and validity of the Chinese version of the social anxiety scale for adolescents. Child Psychiatry and Human Development, 39(2), 185–200.

