## supplement for "Context and Domain Matter: The Error-Related Negativity in Peer Presence Predicts Fear of Negative Evaluation, not Global Social Anxiety, in Adolescents"

**Results of the global social anxiety model**

As described in the main text, AIC model selection revealed that the FNE model provided a better fit to the data. Given that the FNE model exhibited superior fit, subsequent analyses within the main text focus on interpretation of the results of the FNE model. However, for completeness, results of the global social anxiety model are reported below.

Results of the LME model assessing fixed effects of condition (nonsocial vs. social), global social anxiety, and their interaction, on trial-level ERN amplitudes are reported in Table S1. The model found a significant main effect of condition ($t(1317) = -2.668$, $p = 0.008$). However, no significant main effect of global social anxiety, nor an interaction between global social anxiety and condition were identified.

Table S1. **Model 3:** ERN ~ Conditions * GSA

| Predictors | β [95%CI] | SE | t | p |
|---|---|---|---|---|
| (Intercept) | -0.458 [-0.619, -0.296] | 0.083 | -5.545 | < .001*** |
| Conditions | -0.069 [-0.119, -0.018] | 0.026 | -2.668 | .008** |
| GSA | -0.151 [-0.315, 0.012] | 0.079 | -1.925 | .068 |
| Conditions*GSA | -0.029 [-0.080, 0.021] | 0.026 | -1.139 | .255 |

Observations = 1342, N = 23. GSA = Global social anxiety.
*Note*: * $p < .05$, ** $p < .01$, *** $p < .001$.

**Results of the FNE model while controlling for global social anxiety**

Within the main text, we report that the FNE model revealed a significant condition-by-FNE interaction, such that higher levels of FNE were more strongly associated with a larger (more negative) ERN in the social (compared to nonsocial) condition. To determine whether this effect would remain significant when also controlling for global social anxiety, we carried out a supplemental analysis in which global social anxiety symptoms were added to the FNE model as a control variable. Specifically, an LME model assessing fixed effects of condition (nonsocial vs. social), FNE, and their interaction, while controlling for global social anxiety, was conducted (see Table S2). Crucially, this model again revealed a significant condition-by-FNE interaction ($\beta$ = -0.052, 95%CI [-0.102, -0.002], *SE* = 0.026, $p$ = .042) while controlling for global social anxiety. This model demonstrates that FNE predicts unique variance in ERN, while controlling for global social anxiety symptoms.

Table S2. **Model 4:** ERN ~ GSA + Conditions * FNE

| Predictors | β [95%CI] | SE | t | p |
|---|---|---|---|---|
| (Intercept) | -0.462 [-0.631, -0.292] | 0.087 | -5.326 | < .001*** |
| GSA | -0.093 [-0.311, 0.126] | 0.105 | -0.885 | .387 |
| Condition | -0.068 [-0.119, -0.018] | 0.026 | -2.658 | .008** |
| FNE | -0.102 [-0.336, 0.132] | 0.112 | -0.910 | .374 |
| Conditions*FNE | -0.052 [-0.102, -0.002] | 0.026 | -2.033 | .042* |

Observations = 1342, N = 23. FNE = Fear of Negative Evaluation, GSA = Global social anxiety.
*Note*: * $p$ < .05, ** $p$ < .01, *** $p$ < .001.
